# Dolutegravir Developmental Toxicity is Mitigated by Magnesium and Folate in Zebrafish Embryos

**DOI:** 10.1101/2025.06.24.661395

**Authors:** Robert M Cabrera, Ahmed Mohamed, Ryoko Minowa, Katheryn A Neugebauer, Daniel A Gorelick

## Abstract

Integrase-strand-transfer inhibitors have transformed HIV therapy, yet the widely prescribed drug dolutegravir (DTG) has been linked to developmental toxicity and its teratogenic mechanism remains uncertain. Here we use zebrafish to dissect DTG toxicity during early vertebrate development. DTG exposure from 2–4 h post-fertilization (hpf) to 24 hpf produced high mortality and abnormal morphology; co-treatment with folic acid or 5-methyltetrahydrofolate partially restored normal morphology, whereas calcium had no effect. Strikingly, supplementation with magnesium (Mg) rescued DTG-exposed embryos almost as effectively as folate, implicating magnesium availability in protection. In competitive binding assays, Mg increased binding of folate to purified folate receptor (FOLR1) by 30% in the presence of DTG. Maternal-zygotic zebrafish *folr1* mutant embryos contained significantly less endogenous folate than wild-type embryos yet displayed marked hypersensitivity to DTG that could not be mitigated by folate supplementation. Critically, magnesium supplementation partially rescued DTG toxicity in *folr1* mutants, indicating a Folr1-independent component and placing the free DTG vs Mg-bound DTG balance upstream of folate transport. These results support a model in which free DTG antagonizes FOLR1, and Mg modifies DTG developmental toxicity by both FOLR1 dependent and independent processes. Our work identifies magnesium status as a modifiable determinant of DTG teratogenicity and provides a proof-of-concept zebrafish model that could be adapted for rapid *in vivo* screening of integrase inhibitors for developmental phenotypes.

## INTRODUCTION

Dolutegravir (DTG) is an effective and preferred antiretroviral therapy worldwide owing to its potency, high genetic barrier to resistance and convenient once-daily dosing. In 2018, however, nationwide surveillance in Botswana reported higher prevalence of neural-tube defects (NTDs) among infants whose mothers conceived while taking DTG than among those receiving other regimens or no antiretrovirals (Zash et al., 2018; Zash et al., 2019). More recent analyses have tempered that concern. An Eswatini cohort, conducted in a setting that, like Botswana, lacks mandatory folic-acid fortification, found no excess NTDs or other major malformations in more than 4600 peri-conception DTG exposures (Gill et al., 2023). A U.S. study of medical claims covering 14 million pregnancies detected no additional NTD risk in more than 1,000 peri-conception DTG exposures within a population protected by long-standing folic-acid fortification and widespread prenatal supplementation (Kourtis et al., 2023). The same analysis, however, reported a significantly higher rate of pregnancy loss in women who conceived while receiving DTG and, to a lesser extent, other antiretrovirals, suggesting that early embryonic viability remains a potential vulnerability even in the absence of overt malformations. More recently, the same Eswatini network reported that adverse birth outcomes, including pregnancy loss, were elevated in women with HIV compared with HIV-negative women, yet were not linked to DTG or other antiretroviral classes (Gill et al., 2025). Loss of pregnancy is consistent with overt developmental toxicity observed in animal models exposed to DTG during early development. Together these data suggest that maternal HIV infection and underlying nutritional milieu, rather than DTG alone, drive much of the residual risk. Even so, the biological capacity of DTG to perturb embryogenesis remains unclear, and a mechanistic framework is essential to understand and mitigate risks. Rare teratogenic effects can elude even large epidemiologic datasets unless high-risk subgroups are prospectively identified; folate and magnesium status vary widely across regions and could modulate hazard; and all current and next-generation integrase-strand-transfer inhibitors (INSTIs) share the same metal-chelating pharmacophore, so insights gained here will generalize across the class. Regulatory labelling and clinical counselling should rest on a clear understanding of drug-nutrient interactions.

Two non-exclusive hypotheses dominate current discussion. First, DTG has been shown to act as a partial antagonist of folate receptor 1 (FOLR1) *in vitro* and to trigger folate-responsive malformations in zebrafish embryos (Cabrera et al., 2019). Second, DTG’s antiviral activity depends on chelating the pair of Mg ions in the HIV-1 integrase active site. Analogous Mg sequestration in the embryo could either lower the pool of free magnesium or alter FOLR1 activity, thereby compounding folate stress. Dissecting the interplay between folate transport and magnesium homeostasis is not only clinically relevant, it also promises fresh insight into fundamental requirements for early vertebrate development, where maternal folate loading, transmembrane folate carriers and dynamic Mg gradients remain incompletely understood.

Important knowledge gaps remain: the extent to which magnesium availability modulates DTG-induced teratogenicity *in vivo*, the mechanistic influence of magnesium on DTG–FOLR1 interactions, and the mechanisms by which folate-signaling deficits influence embryonic susceptibility to DTG have yet to be defined.

Zebrafish provide an ideal system to address these gaps. Their externally developing embryos permit precise control of exposure windows, rapid high-content phenotyping, and straightforward genetic manipulation. Importantly, DTG induces malformations in zebrafish only when exposure occurs during the first four hours post-fertilization (Cabrera et al., 2019), a developmental window that parallels the periconceptional period implicated in human epidemiology (Zash et al., 2018), supporting translational relevance. Here, we use this model to (i) quantify DTG toxicity and rescue by folate versus divalent cations in wild-type embryos, (ii) test whether Mg modulates DTG–FOLR1 binding *in vitro*, and (iii) determine DTG sensitivity and Mg rescue potential in maternal-zygotic *folr1* mutants that enter development with folate deficiency.

We show that DTG teratogenicity in zebrafish is mitigated by folate or magnesium supplementation; that magnesium abolishes the antagonism of DTG on purified FOLR1 *in vitro*; and that *folr1* mutants are hypersensitive to DTG yet cannot be rescued by exogenous folate but can be rescued by Mg, revealing a dual mechanism of FOLR1 antagonism and Mg sequestration. These findings identify magnesium status as a modifiable determinant of DTG developmental risk, provide a mechanistic framework for the context-specific human data, and establish a rapid zebrafish platform for screening integrase inhibitors and probing fundamental links between magnesium, folate transport and early vertebrate development.

## METHODS

### Zebrafish

Zebrafish were raised at 28.5°C on a 14 h light/10 h dark cycle in the Baylor College of Medicine (BCM) Zebrafish Research Facility in a recirculating water system (Tecniplast USA). Wild-type zebrafish were the AB strain (Westerfield, 2000) and mutant lines were generated using the AB strain. All procedures were approved by the BCM Institutional Animal Care and Use Committee.

### Embryo collection

Adult zebrafish were allowed to spawn naturally in pairs or in groups. Embryos were collected during 20 minute intervals to ensure precise developmental timing within a group. Embryos were placed in 60 cm^2^ Petri dishes at a density of no more than 100 per dish in E3B media (60×E3B: 17.2g NaCl, 0.76g KCl, 2.9g CaCl_2_-2H_2_O, 2.39g MgSO_4_ dissolved in 1 liter Milli-Q water; diluted to 1× in 9 liter Milli-Q water plus 100 μl 0.02% Methylene Blue) and then stored in an incubator at 28.5°C.

### Chemical exposures

Fertilized embryos were placed in 6 well plates (25 embryos per well) with 3 mL of media per well. Chemical exposure began at 2-4 hpf. Embryos were incubated at 28.5°C in the dark and were assessed for developmental malformations and lethality at 24 hpf by light microscopy. All chemicals were purchased as follows: dolutegravir freebase (AChemBlock catalogue 10313, purity 98%, CAS registry number 1051375-16-6); folic acid (Millipore-Sigma catalogue F8758, purity 97%, CAS 59-30-3), L-5-methyltetrahydrofolate calcium salt (Lianyungang Jinkang Hexin Pharmaceutical Company, Magnafolate Pro, purity 99%, CAS 151533-22-1). Stocks were made in 100% DMSO at 1000× and diluted in E3B embryo media to the final concentration (1×) at the time of treatment. All vehicle controls are 0.1% DMSO. To test the effect of Mg and Ca concentrations on DTG toxicity, E3B media were prepared as above but with varying Mg or Ca concentrations.

### Generation of *folr1* mutant zebrafish

Chemically synthesized gRNAs for *folr1* and purified Cas9 protein were obtained from Synthego Corporation (Redwood City, California). We generated gRNA targeting the following sequence in *folr1* exon 4: GGCTGATGAGTCGTGGCGCC**GGG** (PAM in bold, GGG). One-cell stage embryos were injected using glass needles pulled on a Sutter Instruments Fleming/Brown Micropipette Puller (model P-97) and a regulated air-pressure micro-injector (Harvard Apparatus, PL1–90). Each embryo was injected with a 1 nl solution containing 1 µl of 10 μM gRNA, 1 µl of 20 μM Cas9 protein, 2 µl of 1.5 M KCl and 1 µl Phenol Red. The volume of the mixture was increased to 10 μl with 1× microinjection buffer [10 mM Tris-HCl, 0.1 mM EDTA (pH 8.0) in nuclease-free H_2_O] and incubated at 37°C for 5-7 min before injection as described previously (Burger et al., 2016). Mixtures were injected into the yolk of each embryo. Injected embryos were raised to adulthood and crossed to wild-type fish (AB strain) to generate F1 embryos. F1 offspring were sequenced via tail fin biopsies to identify mutations predicted to cause loss of functional protein. The *bcm44* allele contains a 1 basepair insertion (bold underlined, GAGTCGTGGC<u>**C**</u>GCCGGGAGCG) predicted to cause a missense mutation at amino acid 116 followed by a premature termination codon after amino acid 131.

### Genomic DNA isolation

Individual embryos or tail biopsies from individual adults were placed in 100 μl ELB [10 mM Tris (pH 8.3), 50 mM KCl, 0.3% Tween 20] with 1 μl proteinase K (800 U/ml, NEB) in 96-well plates, one sample per well. Samples were incubated at 55°C for 2 h (embryos) or 8 h (tail clips) to extract genomic DNA. To inactivate Proteinase K, plates were incubated at 98°C for 10 min and stored at −20°C.

### Genotyping

PCR and melting curve analysis was performed as described previously (Parant et al., 2009). PCR reactions contained 1 μl of LC Green Plus Melting Dye (BioFire Diagnostics), 1 μl of Ex Taq Buffer, 0.8 μl of dNTP Mixture (2.5 mM each), 1 μl of each primer [5 μM], 0.05 μl of Ex Taq (Takara Bio), 1 μl of genomic DNA and deionized water up to 10 μl. PCR was performed in a Bio-Rad CFX96 thermal cycler, using black and white 96-well plates (Bio-Rad HSP9665). The PCR reaction protocol was 98°C for 1 min, then 34 cycles of 98°C for 10 s, 60°C for 20 s and 72°C for 20 s, followed by 72°C for 1 min. After the final step, the plate was heated to 95°C for 20 s and then rapidly cooled to 4°C. Melting curves were generated with a Bio-Rad CFX96 Real-Time System over a 70-95°C range and analyzed with Bio-Rad CFX Manager 3.1 software. All mutations were confirmed by sequencing. PCR primers to generate amplicons for HRM: ACTAATGTGTGTGTTCAGGCTG, CATACCAGCTCTCACAGTCC. PCR primers to generate amplicons for sequencing: CTGATTCCTTCAGCATGTGTGG, TGCATGCTGGGAACAAGACA.

### Live imaging

Embryos were imaged with a Nikon SMZ25 microscope equipped with a Hamamatsu ORCA-Flash4.0 digital CMOS camera. Images were equally adjusted for brightness and contrast in Adobe Photoshop Creative Cloud.

### FOLR binding assay

Folate receptor binding studies were performed as previously described (Cabrera et al., 2008; Tukeman et al., 2024). Briefly, bovine folate receptor (Sigma) at a concentration of 50 μg/mL was deposited onto microtiter plates (Corning 3361) using a 2×2 grid printed in a 100 nL of 1x phosphate buffer saline via a robotic liquid handler (iDOT). Plates were sealed with adhesive tape and incubated overnight at 4°C in a cold room. The following day (∼16hrs), plates were returned to room temperature and adhesive tape removed to begin processing. Plates were washed 3x with 1X Tris buffered saline + tween 20 (1x TBST, pH8.0) (Sigma) prior to sample application. Samples were prepared by adding 1% ascorbic acid to 1x TBST (pH∼4), prepared as a 10x concentrate, and added to fish in water to 1X, sonicating 50 fish embryos per sample for 30 minutes, followed by manual pipetting. Samples were heat denatured for 5 minutes at 95°C, spun 10Xg for seven minutes to pellet and supernatant collected for folate determination. Supernatant with neutralized with 5% NaOH, ∼2uL, to ∼pH7.5, then 1:10 FA-HRP (Vitros) was added to the neutralized supernatant. Samples were incubated for 30 minutes on washed plates, washed 3x after incubation, and then imaged with ELISA substrate (SuperSignal ELISA Femto Substrate, Thermo). Plates were imaged on a Quansys imager and data extracted and analyzed using 4-plex grids for the 2×2 arrays.

## RESULTS

### DTG causes embryonic defects in zebrafish that are rescued by folate and magnesium

We exposed wild-type zebrafish embryos to 100 μM DTG or vehicle beginning at 2-4 hours post fertilization (hpf) and assayed morphologic abnormalities at 24 hpf. Compared to vehicle controls, embryos exposed to DTG displayed developmental toxicity, including increased mortality and morphological abnormalities, at 1 dpf (Figure 1A), consistent with previous results (Cabrera et al., 2019). Co-treatment with 600 ng/mL folic acid (FA) or 6 μg/mL 5-methyltetrahydrofolate (5-meTHF) significantly reduced these DTG-induced defects (FA: vehicle mean 94% ± 5% SD normal embryos, DTG 34% ± 25, DTG + FA 65% ± 21, n=10 clutches, 25 embryos per clutch; 5meTHF: vehicle control 97% ± 3.5% normal embryos, DTG 77% ± 3, DTG + 5meTHF 93% ± 4, n=5 clutches, 25 embryos per clutch). Magnesium (Mg, 25 mM) supplementation also partially rescued the developmental outcomes of DTG-exposed embryos (Figure 1B), whereas calcium (Ca) had no protective effect on zebrafish developmental toxicity (Mg experiment: DTG in normal media with 0.3 mM Mg 31% ± 20% normal embryos, DTG in media without Mg 38% ± 17%, DTG in media with 25 mM Mg 61% ± 18%, vehicle control 25 mM Mg 91% ± 8%; Ca experiment: DTG in normal media with 0.3 mM Ca 47% ± 28% normal embryos, DTG in media with 3.3 mM Ca 44% ± 21%, vehicle control 3.3 mM Ca 91% ± 8%). These results indicate that the teratogenic effects of DTG are folate responsive and can be mitigated by increasing available folate or increasing magnesium concentrations, consistent with the hypothesis that the partial antagonism of FOLR1 by DTG is mitigated by Mg.

**Figure 1.**
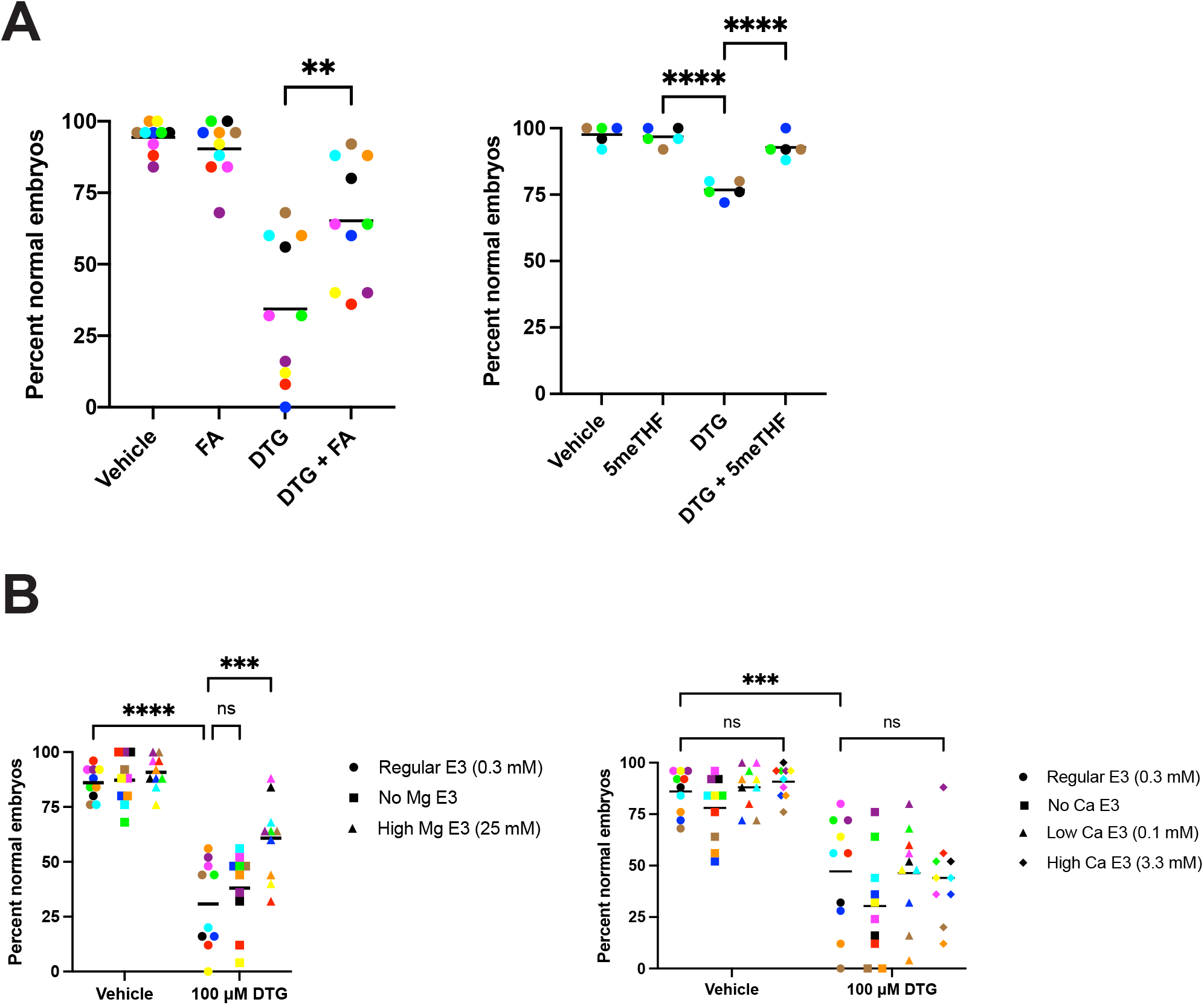
DTG toxicity is rescued by folates and magnesium but not calcium. (A) Wild-type zebrafish embryos exposed to 100 μM dolutegravir freebase (DTG) beginning at 2-4 hpf displayed developmental toxicity, including increased mortality and morphological abnormalities, at 1 dpf. Co-treatment with folic acid (FA, 600 ng/mL) or 5-methyltetrahydrofolate (5meTHF, 6 μg/mL) significantly reduced DTG-induced defects. One-way ANOVA with Tukey’s multiple comparisons test. (B) Magnesium (Mg) supplementation (regular = 0.3 mM, high = 25 mM) partially rescued developmental toxicity of DTG-exposed embryos, whereas calcium (Ca) had no protective effect. Each data point represents mean percent normal embryos from a single clutch of 25 embryos. Within each graph, data points of the same color are from the same clutch. Two-way ANOVA with Tukey’s multiple comparisons test. **, p<0.002; ***, p<0.0002; ****, p<0.0001; ns, not significant.

### Magnesium interferes with DTG–FOLR1 binding in vitro

Competitive folate binding assays were conducted to measure the interactions between folate receptor protein (bFOLR1) and folic acid in the presence and absence of DTG with varying concentrations of Mg. Previous studies indicate that DTG acts as a non-competitive antagonist of FOLR1 at therapeutic concentrations (e.g., 56% signal at 16 μM) (Cabrera et al., 2019). In addition, Mg supplementation (e.g., high Mg diets) reduced the incidence of neural tube defects (NTD) in DTG-exposed mice (Gelineau-van Waes et al., 2023). We found that, in the presence of 100 μM DTG, the interaction between folate and bFOLR1 was increased in the presence of 30 mM Mg versus 0.3 mM Mg (Figure 2; Wilcoxon signed-rank test, p =.008, effect size r = 0.9). These *in vitro* binding assays indicate that Mg modifies the allosteric inhibition of folate receptor by DTG. We conclude that Mg increases folate binding to bFOLR1 in the presence of DTG, supporting the hypothesis that Mg blocks the antagonistic effects of DTG at FOLR1.

**Figure 2.**
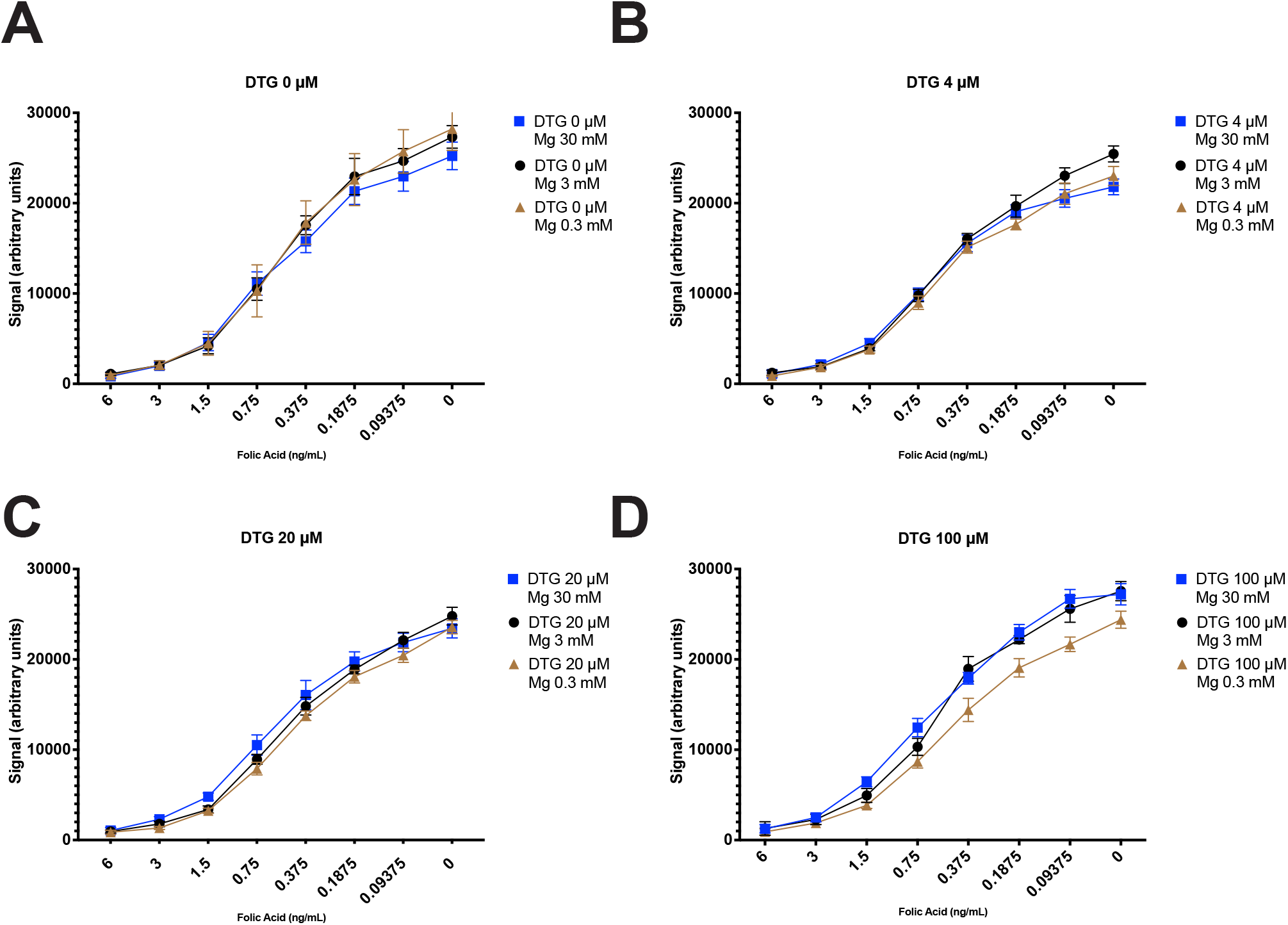
Magnesium increases folate binding to folate receptor in the presence of dolutegravir. In vitro measurements of folate binding to purified bFOLR1 protein in the presence of (A) no DTG, (B) 4 μM, (C) 20 μM, (D) 100 μM DTG together with 0.3 - 30 mM Mg. Folate binding is shown as normalized signal intensity (Y axis). Cold folic acid (X axis) reduces signal. At 100 μM DTG (D), folate binding to bFOLR1 increases with increased Mg concentrations. n=8, mean ± SD.

### Folate receptor mutant embryos exhibit heightened sensitivity to DTG and are not rescued by folate

To directly test the role of the folate receptor in modulating DTG toxicity, we utilized zebrafish carrying a maternal-zygotic mutation in the folate receptor 1 gene (*folr1*) predicted to cause loss of functional protein. In the absence of DTG, MZ*folr1*^*bcm44*^ mutant embryos appeared grossly normal through 1 dpf, though developmental delay was noted with more several morphologic defects at 2 dpf and older (see below). Upon exposure to DTG (100 μM) at 2-4 hpf, MZ*folr1* mutants showed significantly greater developmental toxicity at 24 hpf compared to wild-type embryos at equivalent DTG concentrations (Figure 3A; wild-type vehicle 97% ± 3% normal embryos, MZ*folr1*^*bcm44*^ vehicle 97% ± 7%, wild-type DTG 66% ± 7%, MZ*folr1*^*bcm44*^ DTG 27% ± 19%, n=5 clutches). Whereas DTG-induced developmental toxicity in wild-type embryos could be rescued by co-treatment with FA (600 ng/mL) or 5-meTHF (6 μg/mL), folate supplementation had minimal to no protective effect in MZ*folr1* mutants exposed to DTG (Figure 3B; MZ*folr1*^*bcm44*^ FA experiments: vehicle 97% ± 7% normal embryos, FA 93% ± 6%, DTG 27% ± 19%, DTG + FA 24% ± 10%, n=5 clutches; 5-meTHF experiments: vehicle 99% ± 2%, 5-meTHF 100% ± 0%, DTG 14% ± 26%, DTG + 5-meTHF 10% ± 17%). These findings indicate that a functional folate receptor is required for the rescue effect of exogenous folate and that loss of FOLR1 exacerbates the developmental toxicity of DTG. The inability to rescue folate receptor mutants with supplemental folate further supports that the teratogenic effects of DTG occurs via blocking folate-FOLR1 interactions (rather than downstream folate metabolism).

**Figure 3.**
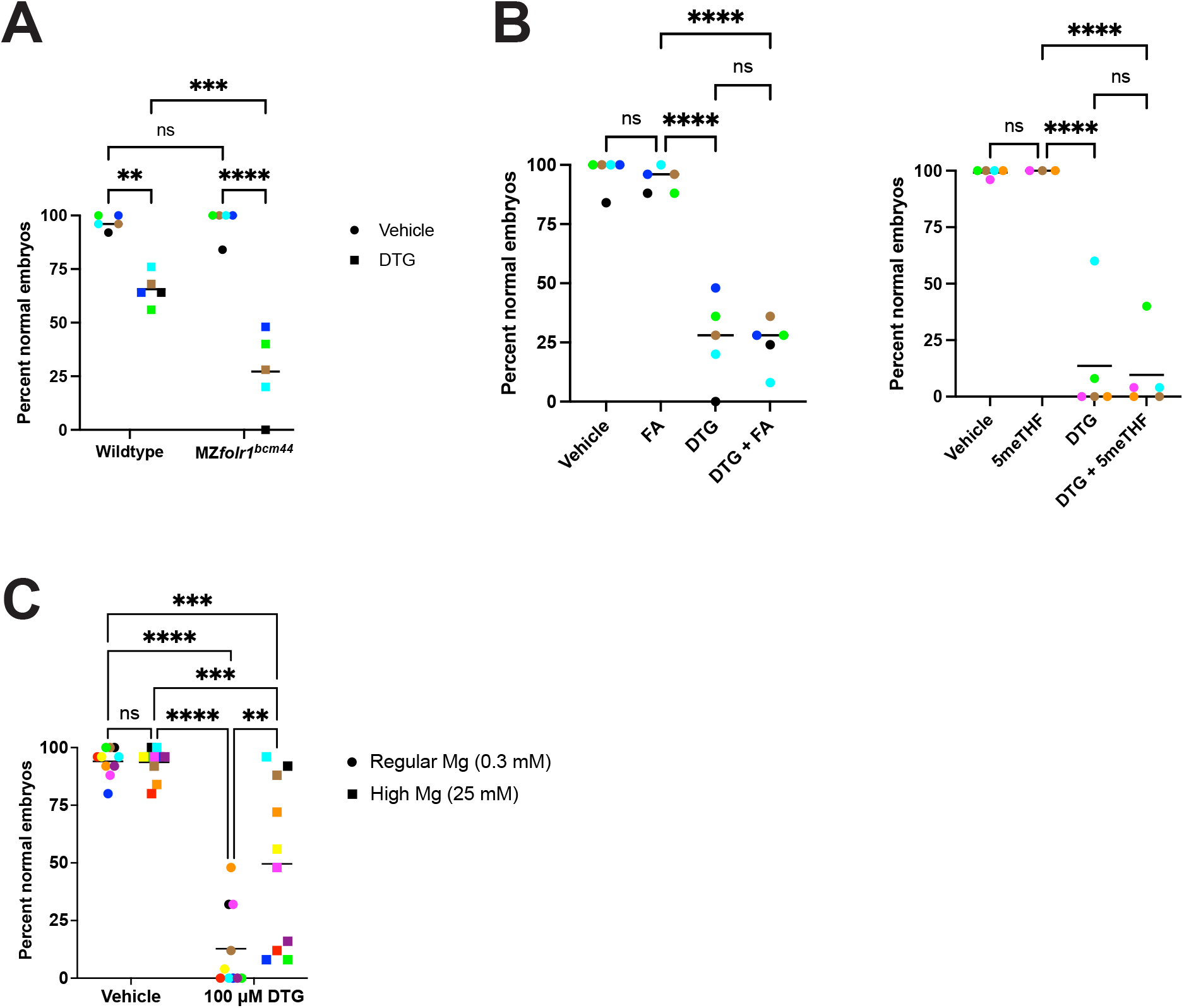
Folate receptor mutants exhibit heightened sensitivity to DTG and are not rescued by folates. To directly test the role of the folate receptor in modulating DTG toxicity, we utilized zebrafish carrying a maternal-zygotic mutation in *folr1* predicted to cause loss of functional protein (MZ*folr1*^*bcm44*^). (A) In the absence of DTG, MZ*folr1* mutant embryos appeared grossly normal through 1 dpf. Upon exposure to 100 μM DTG beginning 2-4 hpf, MZ*folr1* mutants showed significantly greater developmental toxicity compared to wild-type embryos at 1 dpf. Two-way ANOVA with Tukey’s multiple comparisons test. (B) Co-treatment with folic acid (FA, 600 ng/mL) or 5-methyltetrahydrofolate (5meTHF, 6 μg/mL) failed to rescue DTG toxicity in MZ*folr1* mutants. One-way ANOVA with Tukey’s multiple comparisons test. (C) High Mg partially rescues DTG toxicity in MZ*folr1* embryos. Two-way ANOVA with Tukey’s multiple comparisons test. Each data point represents mean percent normal embryos from a single clutch of 25 embryos. Within each graph, data points of the same color are from the same clutch. **, p<0.002; ***, p<0.0002; ****, p<0.0001; ns, not significant.

### Magnesium partially rescues DTG toxicity in folate receptor mutant embryos

Given that exogenous folate cannot rescue *folr1* mutants, we tested whether magnesium modifies DTG toxicity in *folr1* mutants. High Mg (25 mM) partially rescued DTG-exposed MZ*folr1* embryos (Figure 3C): vehicle 94% ± 6% normal embryos; 100 µM DTG 13% ± 18%; vehicle + Mg 94% ± 7%; 100 µM DTG + Mg 50% ± 36%. The improvement from DTG to DTG + Mg was statistically significant (p<0.002, two-way ANOVA), indicating that Mg mitigates DTG developmental toxicity in the absence of Folr1 function. These data place the free (unchelated) DTG vs Mg-bound DTG balance upstream of folate transport and reveal a Folr1-independent component of DTG embryotoxicity.

### Folate receptor mutant embryos have reduced endogenous folate levels and developmental abnormalities

We measured endogenous folate content in MZ*folr1*^*bcm44*^ oocytes and in MZ*folr1*^*bcm44*^ embryos from fertilization through 5 dpf (derived from crosses between *folr1*^*bcm44*^ homozoygous males and *folr1*^*bcm44*^ homozygous females) to assess how loss of the folate receptor affects folate status. Folate quantification revealed that MZ*folr1* mutants possess significantly lower total folate levels in oocytes and embryos compared to wild-type, consistent with impaired maternal folate uptake during oogenesis and early development (Figure 4A). MZ*folr1* mutants, even without any drug exposure, display phenotypes such as smaller head size and cardiac edema beginning at 2 dpf relative to wild-type controls (Figure 4B). These phenotypic abnormalities mirror aspects of folate deficiency in mammalian embryogenesis and suggest that the folate receptor is essential for accumulating sufficient folates for normal development in zebrafish. The folate-deficient state of MZ*folr1* embryos likely predisposes them to additional stressors, explaining their increased susceptibility to DTG at 1 dpf. To confirm that these *folr1* phenotypes are maternally derived, we crossed adult homozygous *folr*^*bcm44*^ females to wild-type males, and wild-type females to homozygous *folr*^*bcm44*^ males, and assayed morphologic abnormalities in embryos through 5 dpf. We found that embryos derived from homozygous mothers exhibited phenotypes similar to MZ*folr*^*bcm44*^ embryos, whereas embryos derived from homozygous fathers appeared normal (Figure 4C). This result supports the idea that female zebrafish deposit folates into oocytes, which are critical for normal embryonic development. Together, our results provide a genetic validation that, in zebrafish, folate receptor function (and by extension folate transport) is crucial for normal development. Additionally, we conclude that folate-FOLR1 interactions provide resilience against DTG-induced developmental toxicity.

**Figure 4.**
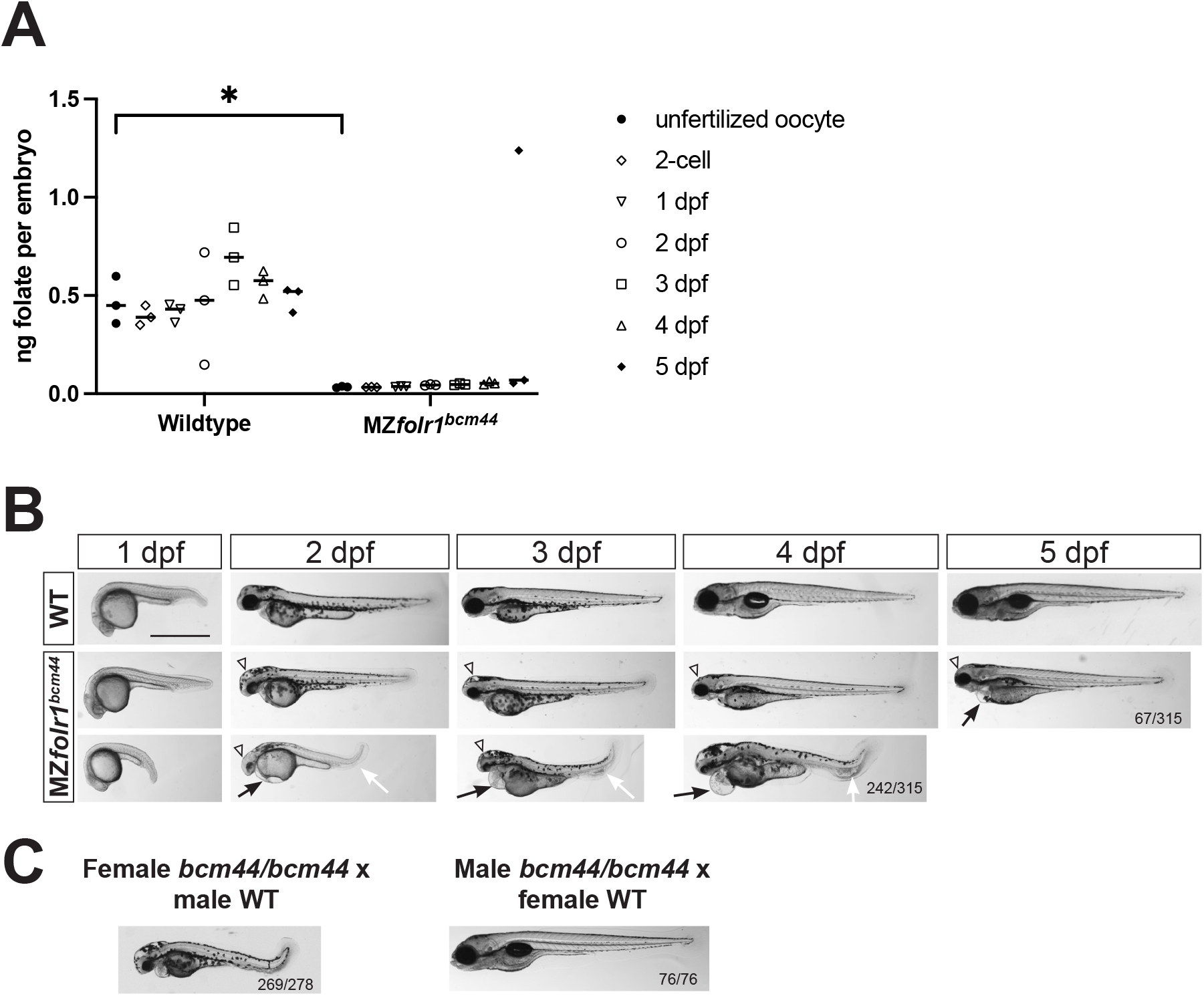
Folate receptor mutant embryos have reduced endogenous folate levels and developmental abnormalities. **(A)** We measured endogenous folate content in MZ*folr1*^*bcm44*^ oocytes and embryos from fertilization through 5 days post fertilization (dpf) to assess how loss of the folate receptor affects folate levels. MZ*folr1* mutants possess significantly lower total folate levels in oocytes and embryos compared to wild-type siblings, consistent with impaired maternal folate uptake during oogenesis and early development. Each dot is folate levels from a pool of 50 embryos. Two-way ANOVA with Tukey’s multiple comparisons test. *, p<0.05. **(B)** MZ*folr1* mutants, without any drug exposure, display phenotypes such as smaller head (arrowheads), cardiac edema (black arrows), and curved tails with ventral thickening (white arrows) through 5 dpf. Examples of mutant phenotypes of varying severity are shown (middle and bottom rows). Scale bar = 1 mm. WT, wild-type embryos. **(C)** *folr1* mutant phenotypes are maternally derived. Female homozygotes crossed to wild-type males produce embryos with phenotypes, whereas wild-type females crossed to homozygous males produce normal embryos. Lateral views with anterior to the left, dorsal to the top. Fractions in the bottom right corners refer to the number of embryos with the indicated phenotype over the total number of embryos examined from 4 clutches (B and C, female *bcm44* x WT) or from 2 clutches (C, male *bcm44* x WT).

## DISCUSSION

### DTG chelates Mg and impairs folate receptor function, leading to folate insufficiency and developmental toxicity

Integrating our findings, we propose a mechanistic model where the teratogenic effects of DTG stem from its magnesium-chelating activity coupled with partial antagonism of the folate receptor. DTG can bind to Mg ions, which not only is the basis for its antiviral action—chelating Mg ions in the active site of HIV-1 integrase, thus blocking the reaction that would splice viral DNA into the host genome (DeAnda et al., 2013; Hare et al., 2011; Passos et al., 2020)—but also chelating and sequestering Mg required for embryonic development (Gelineau-van Waes et al., 2023; Song et al., 2015). By chelating Mg, DTG alters folate receptor interactions as indicated by altered folate binding, effectively reducing folate uptake into the developing embryo. This results in a functional folate deficiency during critical stages of development, causing embryonic phenotypes. Supplemental folate (FA or 5-meTHF) can overcome this blockade in wild-type embryos by mass action, flooding the system with folate to utilize any remaining uptake capacity. Likewise, extra Mg can saturate the chelation capacity of DTG, preserving folate receptor function and normal Mg-dependent cellular processes, thereby rescuing the embryo from the toxic effects of DTG. In *folr1* mutants, exogenous folate cannot rescue because transport via Folr1 is reduced or absent. However, high Mg partially rescued DTG toxicity in *folr1* mutants. This indicates a Folr1-independent component of DTG toxicity and supports a model in which the balance between free DTG and Mg-bound DTG acts upstream of folate transport. Magnesium binds free DTG, producing a chelated form with limited solubility, bioavailability, and affinity for FOLR1 and simultaneously supports Mg-dependent (Mg > DTG) cellular processes.

### Why Mg, but not Ca, protects embryos from DTG toxicity

Our findings that Mg, but not Ca, protect zebrafish embryos from DTG toxicity are supported by biochemical and structural studies of DTG, which show that DTG binds Mg more effectively than Ca. We lack structures of DTG bound to FOLR1, however we can infer DTG preference for Mg over Ca from structures of DTG bound to HIV-1 integrase. DTG chelates two divalent cations, like the twin Mg ions in the integrase active site. High-resolution structures of DTG bound to integrase and viral DNA show DTG binding both catalytic Mg ions. Attempts to substitute Ca abolish DNA strand-transfer activity because the larger Ca ion cannot fit the pocket or achieve the geometry (Hare et al., 2010; Hare et al., 2011; Passos et al., 2020). In free solution the preference is echoed. Pharmacokinetic studies showed that Mg forms a soluble Mg-DTG complex, whereas Ca tends to generate an insoluble precipitate that lowers oral DTG exposure but does not leave a stable, soluble Ca–DTG species in equilibrium (Song et al., 2015). DTG can interact with Ca, enough to reduce oral bioavailability, but the affinity is far lower than for Mg, and the resulting complex is poorly soluble. By contrast, DTG forms a tight, well-soluble chelate with Mg that is the same species captured in every crystallographic structure of DTG-bound integrase.

### Convergence with cation-chelation toxicology

Independent work on intrathecally or intracranially delivered oligonucleotides (ONs) shows that high local ON concentrations can chelate endogenous divalent cations, producing nonspecific phenotypes (side effects) in rodents and non-human primates (Hernandez et al., 2025; Miller et al., 2024; Moazami et al., 2024). Increasing magnesium in the dosing solution mitigated these acute nonspecific effects without impairing ON uptake or activity. These studies attributed the nonspecific toxicity to cation chelation and showed mitigation by raising the Mg:drug ratio. Although the modality and compartment differ from our zebrafish system (intrathecal or intracranial dosing of the central nervous system vs whole-embryo bath exposure), the shared principle is that providing magnesium in the exposure medium or dosing formulation can pre-emptively convert DTG to DTG-Mg-chelate, reducing DTG-folate-receptor interactions, which aligns with our findings that Mg reduces the ability of DTG to antagonize FOLR1 *in vitro* and rescues embryos *in vivo*.

### Translational relevance and human pregnancy data

The requirement that DTG be present only during the first 4 hpf to elicit toxicity in zebrafish (Cabrera et al., 2019) mirrors the narrow periconceptional neurulation window implicated in the Botswana studies (Zash et al., 2018; Zash et al., 2019). A cohort in Eswatini found no excess structural defects among peri-conception DTG exposures (Gill et al., 2023; Gill et al., 2025), whereas a U.S. claims study—conducted in a folate-fortified setting—also reported no additional NTD risk but did detect a significantly higher rate of pregnancy loss in women who conceived while on DTG and a smaller yet similar increase with other antiretrovirals (Kourtis et al., 2023). Pregnancy loss, stillbirth and low birthweight were likewise elevated in the Eswatini cohort regardless of antiretroviral regimen (Gill et al., 2025). Developmental toxicology classifies embryonic death as one of the four canonical manifestations of deviant development. The 24 hpf mortality we observe in DTG-exposed zebrafish thus provides an experimental analogue of these early human losses, while surviving embryos display the malformations noted in the original Botswana signal. Partial rescue of *folr1* mutants by magnesium underscores that maternal magnesium status can buffer DTG-associated risk even when folate transport via Folr1 is compromised. By defining how Mg and folate modulate DTG toxicity, our work provides a mechanistic framework for interpreting these context-specific human data and for guiding targeted surveillance and supplementation strategies.

### Implications for pre-clinical screening

Our data indicate that early zebrafish embryos offer a tractable, 24-hour read-out of DTG-induced malformations that is sensitive to both folate and magnesium modulation. The combination of external development, optical transparency and small molecule permeability enables high-throughput, low-cost testing of drug analogues in a whole-vertebrate context. With automated imaging and computer-vision scoring, the assay could be scaled to 96-well or 384-well format, allowing side-by-side comparison of current and next-generation integrase-strand-transfer inhibitors (INSTIs) for teratogenic potential. Importantly, the platform captures chemical features that influence folate-receptor antagonism and those that affect magnesium binding, providing a unique opportunity to rank-order compounds by developmental safety margins. A zebrafish screen could serve as an early gate to identify integrase inhibitors with reduced developmental risk and to inform medicinal-chemistry optimization of the INSTI scaffold for preclinical testing.

### Limitations

First, the DTG concentrations used in embryo media (100 µM) exceed typical human plasma levels. Without direct pharmacokinetic measurements in zebrafish, we cannot determine the actual intra-embryonic DTG levels. It is possible that supraphysiologic DTG concentrations in embryo media are needed to achieve physiologic concentrations in embryos, as has been shown to occur with other small molecules like estrogens and androgens (Souder and Gorelick, 2017; Souder and Gorelick, 2018; Zadmajid et al., 2024). Second, our conclusion that magnesium modifies DTG developmental toxicity is based on rescue experiments *in vivo*, including partial rescue in *folr1* mutants, and on FOLR1 binding data *in vitro*. We did not measure intracellular free or total Mg in embryos, so the Mg-dependent protection we propose (conversion of free DTG to Mg-bound DTG chelate and support of Mg-dependent processes by increasing the Mg:DTG ratio) remains inferential. Third, we assessed DTG antagonism only at FOLR1 and did not quantify the effects of DTG on other zebrafish folate transporters (reduced folate carrier SLC19A1, proton-coupled folate transporter SLC46A1, mitochondrial folate transporter SLC25A32). Thus, the relative importance of each transporter in mediating or modifying DTG toxicity is unresolved.

### Conclusions

In summary, our model posits that the developmental toxicity of dolutegravir is driven by a dual mechanism of (i) folate pathway inhibition (via FOLR1 antagonism) and (ii) magnesium sequestration, ultimately impairing essential folate-dependent developmental pathways and other Mg-dependent processes necessary for normal embryonic development. *folr1* mutants are already folate-limited, so they have no metabolic cushion against magnesium sequestration that impairs Mg-dependent processes, making them appear more sensitive to DTG exposure. Our findings support a dual-pathway model in which free DTG antagonizes FOLR1, while the ability of DTG to chelate Mg also perturbs Mg-dependent biology. Partial rescue of MZ*folr1* embryos by Mg demonstrates that magnesium acts upstream and independently of Folr1, highlighting the balance between free DTG and Mg-bound DTG as a key driver of developmental outcome. Our results suggest folate and magnesium status as critical modifiers of DTG teratogenic risk in zebrafish embryos, in agreement with recent findings in mice (Gelineau-van Waes et al., 2023; Tukeman et al., 2024), supporting the idea that ensuring adequate folate and magnesium levels are potential strategies for mitigating DTG-associated birth defects, including in settings of compromised folate transport. Larger cohort studies in diverse populations, particularly in settings with varying folate and mineral intake, would help clarify human risk. Additionally, routine monitoring of plasma folate and systematic evaluation of nutritional interactions in patients on DTG, especially during pregnancy, would be expected to provide actionable clinical guidance and help prevent adverse outcomes.

## ACKNOWLEDGEMENTS

We thank Kelli Kesler, Lauren Pandolfo and the staff of the Baylor College of Medicine zebrafish facility for caring for our zebrafish. This work was supported by NIH grant R01HD100229. Conflicts of interest: RMC is a consultant for Dartox Consulting, LLC, and chief scientific officer of AlphaRose Therapeutics. DAG is editor in chief of Biology Open, published by the nonprofit The Company of Biologists.

